# Classification of Preeclamptic Placental Extracellular Vesicles Using Femtosecond Laser-fabricated Nanoplasmonic Sensors and Machine Learning

**DOI:** 10.1101/2021.12.28.474354

**Authors:** Mohammadrahim Kazemzadeh, Miguel Martinez-Calderon, Song Y. Paek, MoiMoi Lowe, Claude Aguergaray, Weiliang Xu, Lawrence W. Chamley, Neil G.R. Broderick, Colin L. Hisey

## Abstract

Placental extracellular vesicles (EVs) play an essential role in pregnancy by protecting and transporting diverse biomolecules that aid in fetomaternal communication. However, in preeclampsia, they have also been implicated in contributing to disease progression. Despite their potential clinical value, most current technologies cannot provide a rapid and effective means of differentiating between healthy and diseased placental EVs. To address this, we developed a fabrication process called laser-induced nanostructuring of SERS-active thin films (LINST), which produces nanoplasmonic substrates that provide exceptional Raman signal enhancement and allow the biochemical fingerprinting of EVs. After validating LINST performance with chemical standards, we used placental EVs from tissue explant cultures and demonstrated that preeclamptic and normotensive placental EVs have classifiably distinct Raman spectra following the application of both conventional and advanced machine learning algorithms. Given the abundance of placental EVs in maternal circulation, these findings will encourage immediate exploration of surface-enhanced Raman spectroscopy (SERS) as a promising method for preeclampsia liquid biopsies, while our novel fabrication process can provide a versatile and scalable substrate for many other SERS applications.

---

Preeclampsia is a common pregnancy complication which presents as new onset hypertension after mid gestation and can endanger both the mother and baby before and after birth. It is associated with damage to the endothelium and vasoconstriction, which can eventually lead to the dysfunction of various organ systems and is estimated to result in the death of more than 50,000 young women each year.^1^ Preeclampsia can be divided into early or late onset depending on when it first presents during pregnancy, with the early onset variant often being more severe with higher risk of morbidity and mortality to both the mother and baby. However, the actual causes and mechanisms for the progression of preeclampsia are not well understood, and despite extensive research, few biomarkers are known which can predict or monitor the presence of the disease. Extracellular vesicles (EVs), which normally play a vital role in fetomateral communication during healthy pregnancies,^2,3^ are one potential mediator of preeclampsia.^4,5^ In particular, the syncytiotrophoblast is a large, multinucleated cell that covers the entire surface layer of the placenta and is bathed in maternal blood, extruding vast quantities of heterogeneous placental EVs into the maternal circulation.^6^ These syncytiotrophoblast-derived EVs are implicated in the progression of the disease, and also hold promise as biomarkers for the early diagnosis and treatment monitoring of preeclampsia given their abundance and accessibility in the maternal circulation.^4,7^

Conventional characterization approaches have provided valuable information related to the biochemical cargo of EVs in preeclampsia compared to healthy pregnancies, including varying levels PLAP and numerous miRNAs.^8–14^ However, several potential characterization tools remain largely unexplored. One such tool, Raman spectroscopy, has recently shown promise for EV characterization, particularly when paired with nanostructured plasmonic surfaces.^15^ These plasmonic surfaces effectively amplify the normally weak EV Raman signal by several orders of magnitudes through the generation of strong nanoscale electric fields, making it possible to biochemically fingerprint EVs even down to the single EV level.^16^ Following the application of machine learning algorithms, the acquired spectra can then be used to classify different subpopulations of EVs.^17^ The use of Surface Enhanced Raman Spectroscopy (SERS) to classify EVs has been primarily used in cancer applications,^18–20^ but has recently begun to expand into other EV-related research fields such as bacterial or mesenchymal stromal cell (MSC) EV classification.^21,22^ Clearly, the utilization of SERS for EV characterization in preeclampsia or other pregnancy diseases could provide immense value given its label-free and generally nondestructive nature.

However, for EV SERS to become a routine research or clinical characterization method, cost-effective and versatile plasmonic surfaces with high sensitivity must be available. These surfaces should be easy to manufacture, stable over time, and highly reproducible from batch to batch. In the past, two primary methods of SERS substrate fabrication have been utilized, including colloidal suspension and direct nanopatterning of solid surfaces.^23,24^ Colloidal suspensions involve the deposition of a layer of nanoparticles onto a surface, mixture within hybrid materials, or direct mixture with EV suspensions which can then either be used directly or as a template for subsequent replica molding, depending on the approach.^25–31^ Alternatively, many roughening or patterning techniques can be used to create nanostructures on various surfaces, however their fabrication often relies on availability of nanometric precision lithography systems within cleanrooms, such as hole-mask colloidal, focused ion beam, electron beam, or photo-lithography.^22,23,32,33^ Importantly, access to these facilities are often costly and require additional materials which can severely limit accessibility and routine usage outside of laboratory research settings.

In contrast, femtosecond (fs) laser ablation offers a scalable, tunable, single step, and reproducible approach to SERS substrate fabrication.^34–38^ Many efforts have been focused on exploiting fs-laser machining to tune surface wettability, creating combinations of superhydrophobic/philic areas that enhance detection performance by concentrating analyte molecules into hotspot areas, leading to the detection of concentrations down to 10^−13^*M* using SERS.^39–42^ Additionally, fs-laser induced plasma assisted ablation has been used to create active SERS substrates based on Ag nanoparticles, demonstrating promising results regarding food safety detection. ^43^ Other laser-based surface roughening approaches for SERS substrate fabrication mainly leverage the phenomenon known as laser induced periodic surface structures (LIPSS), which allows the fabrication of nanopatterns with sub-wavelength resolution.^44–46^ These efforts have been directed solely towards LIPSS nanostructuring of base materials, such as silicon or glass, followed by the deposition of a SERS-active film of gold or silver.^47–53^

Despite the potential benefits, fs-laser nanopatterning directly on SERS-active thin films has not yet been demonstrated. Although several studies have demonstrated LIPSS patterning of other types of thin films,^54–56^ the fs-laser patterning of SERS-active films has likely been avoided due to the significant risk of completely ablating the somewhat delicate and thin SERS-active layer due to the use of high intensity pulses, even when using the low powers required for LIPSS formation. Directly patterning the SERS-active thin films could provide advantages in terms of fabrication scalability, and could improve the Raman signal enhancement compared to subsurface patterning since coating a base pattern with a SERS-active thin film overlay automatically smooths the surface features, thus reducing the curvature. Furthermore, fs-laser ablation naturally results in the generation of particle debris which often redeposits onto the patterned surface.^57^ For SERS applications, these redeposited gold or silver nanoparticles could theoretically produce even greater Raman signal enhancement than the base nanopattern alone.

In this study, we developed a method known as laser-induced nanostructuring of SERS-active thin films (LINST) and demonstrate that it can produce scalable and reproducible SERS substrates for EV fingerprinting and classification. Instead of patterning the base substrate such as silicon or glass, we show that direct fs-laser machining of gold thin films easily achieves the SERS amplification needed to obtain EV Raman spectra and that this SERS amplification is at least partially due to the redeposition of gold nanoparticle debris. Finally, following the application of conventional and advanced machine learning on the acquired SERS spectra, healthy and preelcamptic EVs from tissue explant cultures are efficiently classified. To the best of our knowledge, this is the first example of using fs-laser-machined nanoplasmonic surfaces, LIPSS or otherwise, for EV SERS applications.

## Results and Discussion

### Formation of LINST

The LINST process enabled the soft nanostructuring of gold thin films, avoiding the complete removal of the gold layer while creating nanoscale roughness across the entire surface that enabled straightforward SERS analysis of both chemical species and EVs. The laser scanning strategy was tailored to very low fluence (slightly above the ablation threshold) and the distance between scanned lines was optimized to continuously pattern the whole area (detailed in methods). Ultimately, the optimized LINST was carried out with a fluence of 0.2 J/cm^2^, scanning speed of 1.125 mm/s, and a separation between the scanned lines of 2.5 *μ*m.

Both magnetron sputter coated and thermal evaporated gold thin films were tested for their potential use in LINST patterning. As shown in Figure 1a-b, the initial grain size was different for each method, as well as the overall thin film morphology at the nanoscale (detailed in methods). However, following fs-laser-machining, the LINST were almost identical in terms of nanotopography (Fig 1c-d), and both types of gold thin films were fully covered by nanostructures with no areas remaining unpatterned. As expected, the initial differences in grain size did not play a major role in the LINST process since the absorption of both surfaces was very similar and both thin films were of similar thickness.

**Figure 1:**
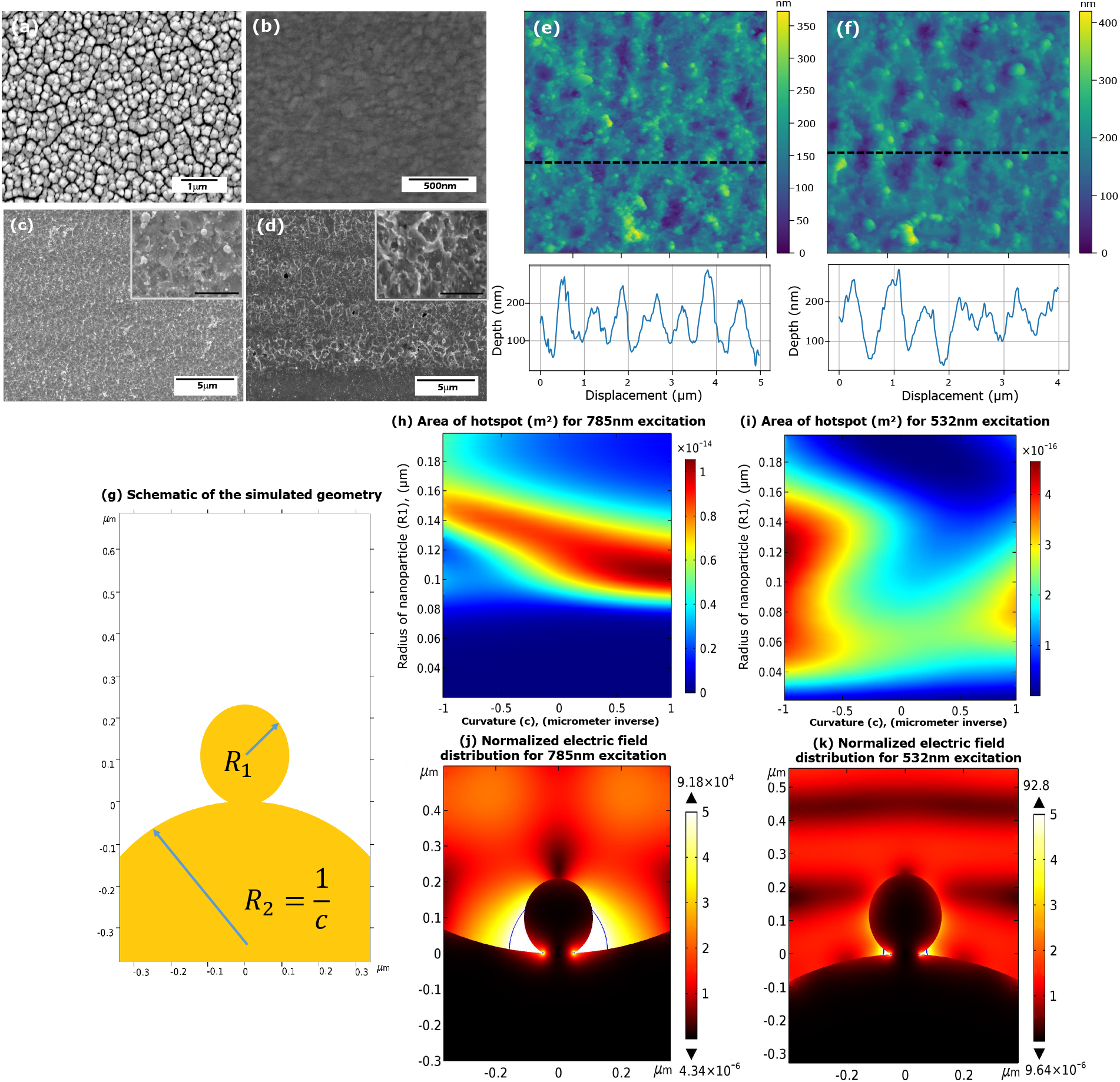
SEM image of the gold thin films and resulting LINST patterns for (a,c) magnetron sputter coated and (b,d) thermal evaporated gold thin films (insets are higher magnification images of the same nanopatterned area, scale bars = 1 *μ*m). AFM images and profiles for LINST on (e) magnetron sputter coated and (f) thermal evaporated gold thin films. COMSOL simulation results of LINST plasmonic surfaces showing (g) a schematic of the simulated geometry, (h) hotspot areas for 785 nm for different substrate curvatures and particle radii, (i) hotspot areas for 532 nm excitation for different substrate curvatures and particle radii, normalized electric field distributions for instances with maximum hotspot areas using (j) 785 nm excitation and (k) 532 nm excitation.

The presence of slightly periodic nanoripples oriented perpendicular to the scanning direction can also be seen, which somewhat resembles the well-known LIPSS. Most recent LIPSS studies conclude that the formation of such structures arise from a combination of plasmonic interference effects and hydrodynamic matter reorganization.^58^ In the case of LINST, the need to nanopattern the surface while completely avoiding bulk removal of the thin film led to the use of a very low fluence and relatively low number of pulses. These conditions made LINST patterns less defined and coherent compared to LIPSS, but the resulting periodicity combined with the gold nanoparticle redeposition ultimately produced effective SERS substrates.

2D-FFT analysis of the SEM images indicated that the fabricated patterns had an average periodicity of 565±20nm and 548*nm*±27nm which is slightly smaller than the irradiation wavelength and in agreement with the expected periodicity of the well known Low Spatial Frequency LIPSS. Additionally, AFM images and profiles (Fig 1e-f) corroborate these findings and show the presence of both convex and concave structures, around 150 nm deep, across the entire patterned surface. Given the known thickness of the deposited gold layer, the AFM results (and later SERS results) verify that the gold thin films were not fully ablated at any point, avoiding any risk of the underlying chromium or silicon substrate contaminating the SERS signal. Both SEM and AFM also clearly show the presence of small gold nanoparticles (tens of nanometers) covering the entire patterned area, which are caused by the redeposition of the ablated material (Fig 1c-d insets). Once it was established that either deposition method could be easily used for LINST patterning, thermal evaporation was used for the remainder of the study for scalability and more consistent purity.

### Numerical Simulations

As seen in Figure 1, the curvature of the gold LINST substrates varies from positive to negative while gold nanoparticles are scattered across the entire surface. In order to estimate the effect of these combined features on the Raman signal enhancement, simulations were performed in COMSOL Multiphysics. In our simulations there are three parameters of interest: *R*_1_ is the radius of the gold nanoparticle, *c* is the local curvature of the surface, and λ is the wavelength of the incident light. We further only simulate a 2D geometry as shown in Fig. 1g which captures the essential features while reducing the computational time down to the point where parameter sweeps over the wide range of possible parameters are possible. This is necessary since the nanoparticle size and substrate curvature are randomly distributed around their mean value and thus a single simulation will not provide an accurate value for the surface as a whole. A small portion (10 nm) of the nanoparticle was also fused into the substrate to avoid a singularity at their intersection^59,60^ and to create more realistic simulations.

As previously established, plasmonic nanostructures can concentrate incoming light near their nanometric features and sharp edges.^21,23^ Such hotspots lead to significantly greater but highly localized electric field amplitudes, effectively enhancing any nonlinear phenomena such as Raman scattering. To quantify this effect for various geometries and conditions, we calculated the area of regions which have an electric field enhancement of more than 5 (corresponding to more than 5^4^ fold Raman enhancement) as shown Figure 1h and 1i for 532 nm and 785 nm excitation wavelengths, respectively. These simulations focused on 532 nm and 785 nm light as these wavelengths are available in our setup and are also the most commonly used wavelengths for Raman spectroscopy, but the same approach could be easily adapted to other excitation wavelengths due to the scale invariance of Maxwell’s equations.

According to our simulations, 785 nm radiation has up to a 100 times larger hotspot area for the particle sizes and substrate curvatures that were tested. This should in turn lead to a better Raman signal enhancement for larger molecules and EVs as they can more easily fit within the hotspots. By contrast, smaller molecules could be analysed with either laser as they can fit into the 532 nm hotspots. Finally Figure 1k and 1j show the normalized electric field relative to the incident field for the optimized structures for both 532 nm and 785 nm, respectively. In these figures the color map stops at a normalized strength of 5, although the maximum is significantly higher in each case.

### SERS Chemical Measurements

To clearly demonstrate the performance of the optimized LINST for SERS of chemical species, Rhodamine 6G (R6G) solutions of concentrations ranging from (10^−5^*M*) to (10^−8^*M*) were deposited and dried on the fabricated samples. Then, the Raman spectra of R6G at each concentration were acquired using a 532 nm laser and 1 second acquisition time with a 50 × microscope objective (NA=0.42). The average of 100 spectra at each concentration and their standard deviation are shown in Supplementary Figure 1. This demonstrates the high sensitivity of the fabricated SERS as all the R6G bands are clearly observable even at a concentration of (10^−8^*M*). There is also a clear quantitative relationship between R6G concentration and the amplitude of the signal, meaning that with proper calibration, it could potentially be used to gain concentration information of the sample from the acquired SERS spectra alone.

#### SERS Spectral Quality Improvement

Hyperspectral Raman imaging was also used to compare the sensitivity of the LINST area to the area which was not directly machined. In this case, the 532 nm laser with 50× microscope objective and 10^−7^*M* R6G were used, but as a relatively large grid (100 × 100 *μ*m) was imaged, the acquisition time was reduced to 100 ms. As shown in the optical image of the investigated sample (Figure 2a), an area which contains the boundary between the machined area (lower half) and bare gold (upper half) was chosen to determine if any correlation between machining and obtained spectra quality existed. In Figure 2b, the amplitude of the obtained spectra at 1358 *cm*^−1^, one of the main R6G peaks, is shown. Although this slightly higher amplitude of the peak of R6G over the machined area shows a better performance of the machined area, the information regarding the other peaks and hence the overall quality of the obtained spectra is not clearly demonstrated.

**Figure 2:**
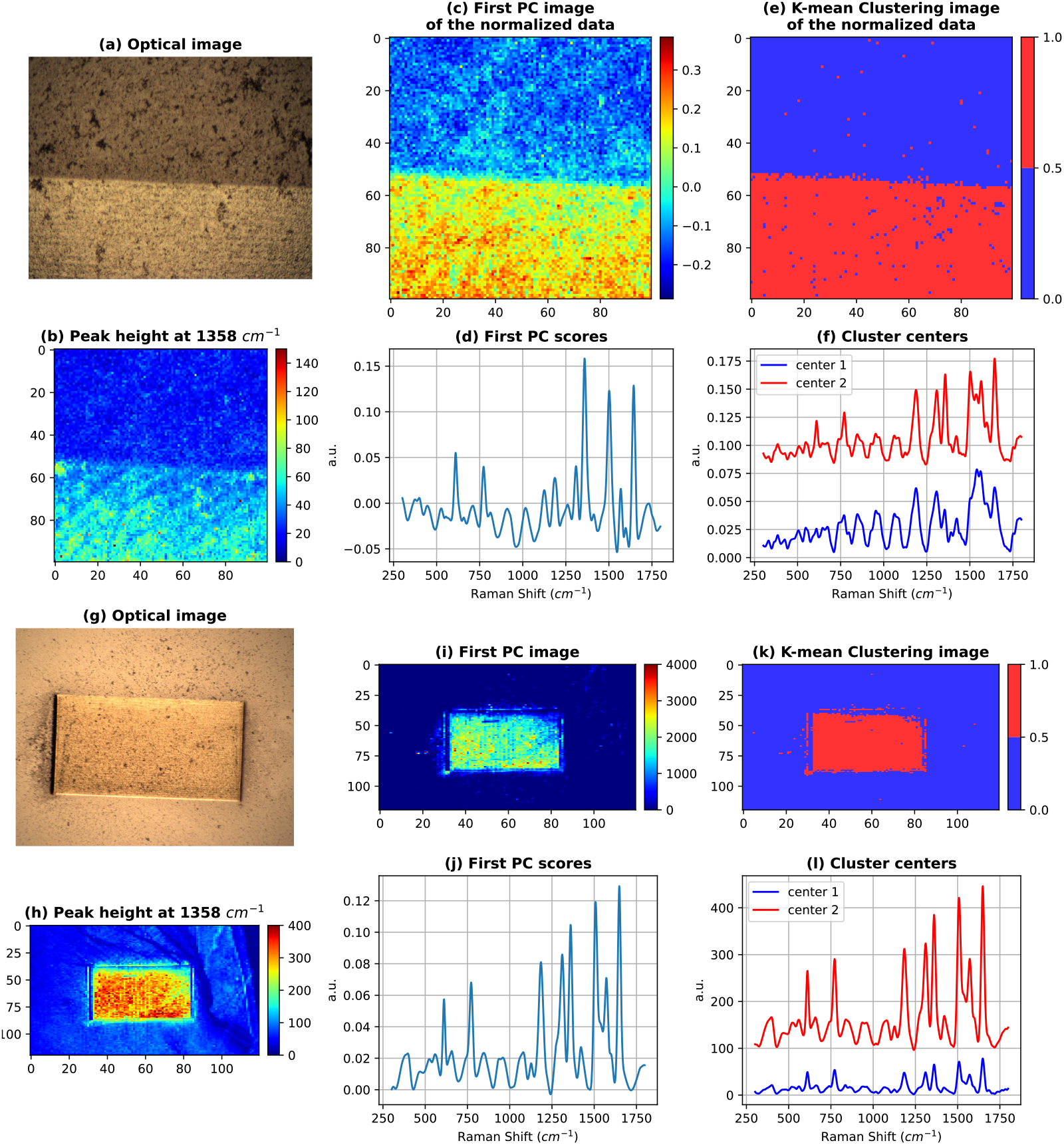
LINST SERS using 10^−7^*M* R6G showing (a) optical image of the Raman imaging area used in (b) Raman image based on peak height at 1358*cm*^−1^, (c) corresponding amplitude of the first PC of the euclidean normalized Raman image spectra, (d) PC score of (c), (e) distribution of 2 clusters obtained using K-mean clustering, and (f) center of the clusters with their corresponding colors in (e). LINST SERS using 10^−6^*M* R6G showing (g) optical image of the larger Raman imaging area used in (h) Raman image based on peak height at 1358*cm*^−1^, (i) amplitude of the first PC, (j) PC score of (i), (k) distribution of 2 clusters obtained using K-mean clustering, and (l) center of the clusters with their corresponding colors in (k).

To better understand the overall quality of the acquired spectra rather than a single peak’s amplitude, the first principal component (PC) was calculated for the normalized spectra and used for imaging as shown in Figure 2c for the first PC score shown in Figure 2d. Interestingly, this PC score clearly contains all the major peaks of R6G and thereby higher intensity of the depicted data in the PC image on machined area in Figure 2 (c) actually demonstrates better signal quality. However, for the PC calculation, all the investigated points in the imaged area play an equal role, distorting the first PC from bare gold which likely has much lower quality.

To compensate for this drawback of PC imaging, K-mean clustering was used to automatically label the normalized and unlabeled data into two clusters, with the center of each cluster being its representative spectrum. The K-mean imaging for R6G when different colors (blue and red) are used to represent the different clusters is shown in Figure 2e. The cluster centers are also shown in Figure 2f for their corresponding colors in Figure 2e. Clearly, the boundary of the machined area corresponds to the boundary of the K-mean clusters of the normalized data. Moreover, the fact that the cluster center corresponding to the machined area more clearly contains all the R6G peaks compared to the cluster center for the bare gold proves that the machined area significantly enhances the quality of the spectra at this concentration. Finally, the scattered points with greater signal quality on the bare gold, but in close promximity to the LINST area (Figure 2e), indicate the presence and Raman enhancement capability of redeposited gold nanoparticles alone.

#### Spectral Amplitude Enhancement

To investigate the amplitude of the obtained spectra, an image of the LINST was taken with the 532 nm laser, 5× microscope object, 25% of ND filter, and R6G concentration of the 10^−6^*M*. As a higher concentration of R6G was used, the bare gold also produced a Raman signal with acceptable quality, and the difference in this example should be solely due to the amplitude of the obtained signal. In this case, the data which are used for the statistical analyses are not normalized to preserve the amplitude in their result. An optical image corresponding to the Raman hyperspectral imaging area, amplitude of the obtained spectra at 1358 *cm*^−1^, first PC image, first PC score, K-mean clustering image, and their corresponding centers are shown in Figure 2g-l, respectively. In this case, the peak at 1358 *cm*^−1^ shows a much stronger signal in the machined area. As seen in Figure 2j, the first PC score is clearly the expected R6G spectrum, and thus any higher amplitude in Figure 2i directly indicates higher intensity of the obtained spectra. Similarly, the K-mean clustering image identified the machined area as a separate cluster and its corresponding center shows how much higher, on average, the signals in the machined area are in comparison to the bare gold surface (Fig 2k-l). Together, these results clearly demonstrate the improvements in signal strength and quality provided by the LINST substrates for R6G, which should also be the case for most other chemical analyses.

### EV SERS Spectra

In a similar approach to the previous R6G testing, hyperspectral Raman imaging of the LINST surface following the release and drying of one of the late onset preeclamptic (LOPE) EV suspensions was performed. The Raman acquisition parameters were: 785 nm laser with a 25% ND filter, 10× microscope object and .3s (300ms) acquisition time. K-mean clustering was again used to automatically label the data based on its amplitude and signal shape. The optical image of the investigated area, K-mean clustering image, and corresponding cluster centers are shown in Figure 3a-c. Clearly, the LINST area produces characteristic EV SERS spectra while the flat gold surface produces little to no signal, with the exception of the LINST-adjacent region containing redeposited gold nanoparticle debris. This EV Raman signal from the regions of flat gold with redeposited nanoparticles is in agreement with the simulations presented in Figure 1h, where regions of zero curvature still create a hotspot for certain nanoparticle sizes when using the 785 nm laser.

**Figure 3:**
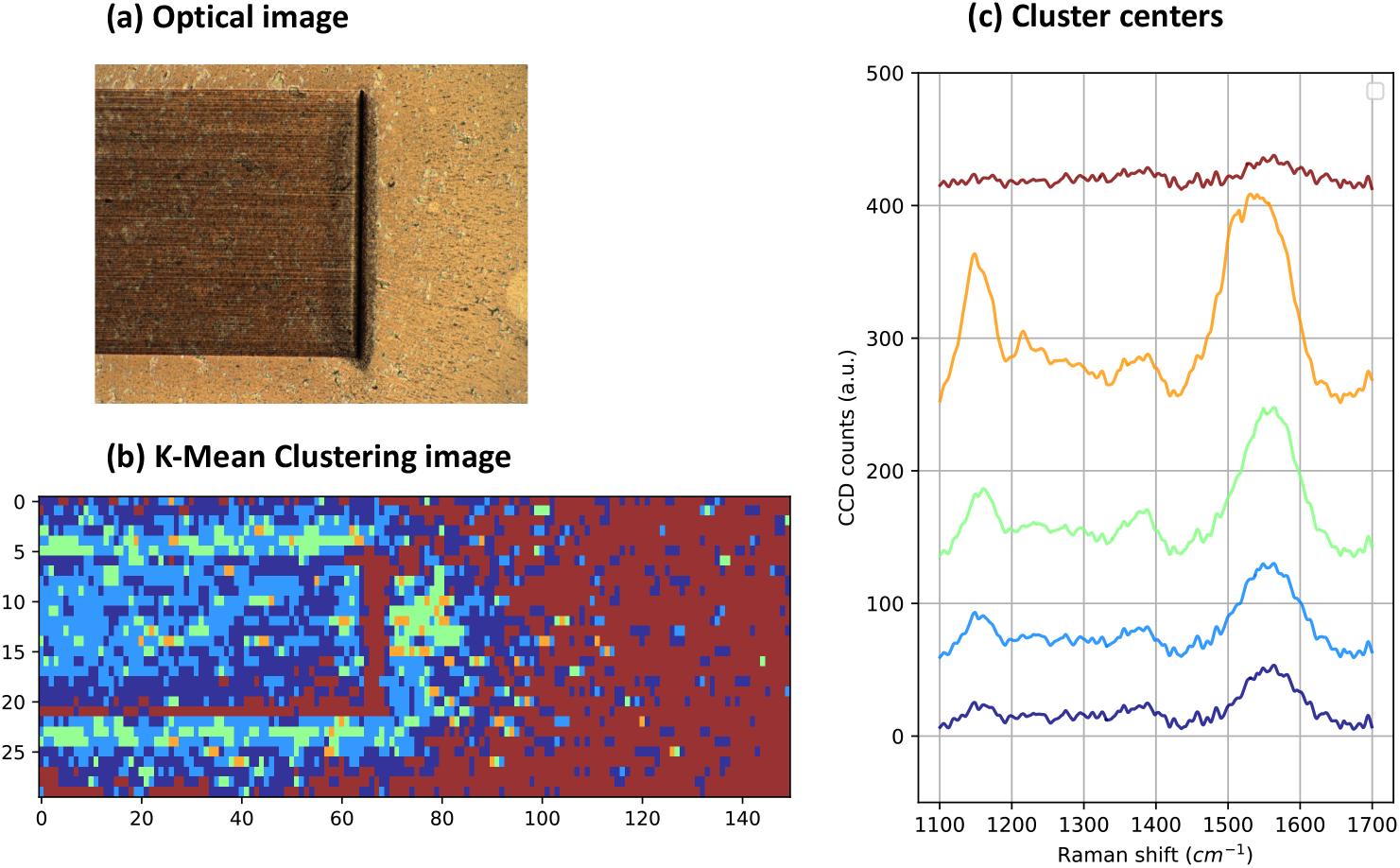
LINST SERS using LOPE EVs showing (a) optical image of area used to obtain the Raman imaging, (b) distribution of 5 K-mean clusters over the imaged area, (c) center of each cluster depicted with their corresponding colors in (b).

This finding is also in agreement with previous finding of EV SERS spectra acquisition using “nanocups” and flat gold surfaces. ^18^ In fact, due to relatively large sizes of small EVs (30-150 nm) compared to chemical species and EVs’ lack of strong chromophore molecules, their SERS spectra are much weaker than those of chemical dyes like R6G.^61^ Their size also prevents them from fitting perfectly into the nanometric hotspots, in theory resulting in larger EVs producing weaker signals compared to the smaller EVs due to their curvature. Thus, only SERS which have larger hotspot areas or are perfectly designed for EV sizes and curvatures are suitable for EV SERS measurements in contrast to flat gold surfaces, which provides weak to no Raman spectra for EVs.

### Classification of Normotensive and Preeclamptic Placental EVs

SERS spectra of small EVs of the same concentration and isolated from 13 different tissue explant cultures, including 5 normotensive (NT), 5 early onset preeclampsia (EOPE), and 3 late onset preeclampsia (LOPE), were acquired (culture and EV isolation detailed in methods). For each sample, 100 spectra between 800*cm*^−1^ to 1800*cm*^−1^ wavenumbers were recorded over a 10×10 *μ*m rectangular grid on the optimized LINST due to the expected heterogeneity. To obtain these spectra, the Raman microscope configuration of a 50 × microscope object, 785 nm laser, 10 s acquisition time, and a 10% ND filter were used.

A conventional approach for reporting EV Raman spectra is shown in Fig. 4, where the normalized spectra are averaged and presented with their standard deviation, removing any information on exact spatial heterogeneity. In Figure 4a,c and e the averaged size distributions from nanoparticle tracking analysis (NTA) show the consistent isolation of small EVs (50-200 nm) from differential ultracentrifugation followed by size exclusion chromatography (SEC), ensuring that only EVs of similar sizes were compared. In addition, in Figure 4b, d and f the averaged spectra more clearly demonstrate some key differences between samples types. Here, the average spectrum is presented as the blue line with standard deviation in yellow, with each subtype of biomolecules peaks indicated by corresponding color bands as noted above the spectra. Of particular interest is the peak at around 1750 *cm*^−1^, which is clearly more prominent in the normotensive samples as well as more subtle differences in the shapes and sizes of some peaks. The 1750 *cm*^−1^ peak is known to be directly related to the stretching C=O double-bond of esters found in lipids and phospholipids, based on past Raman studies of other biological membranes and lipid reference products.^62–64^ Similarly, there appear to be additional peaks at 870 *cm*^−1^ and 1330 *cm*^−1^ in the normotensive samples, which could be caused by several known phospholipid-associated bonds.

**Figure 4:**
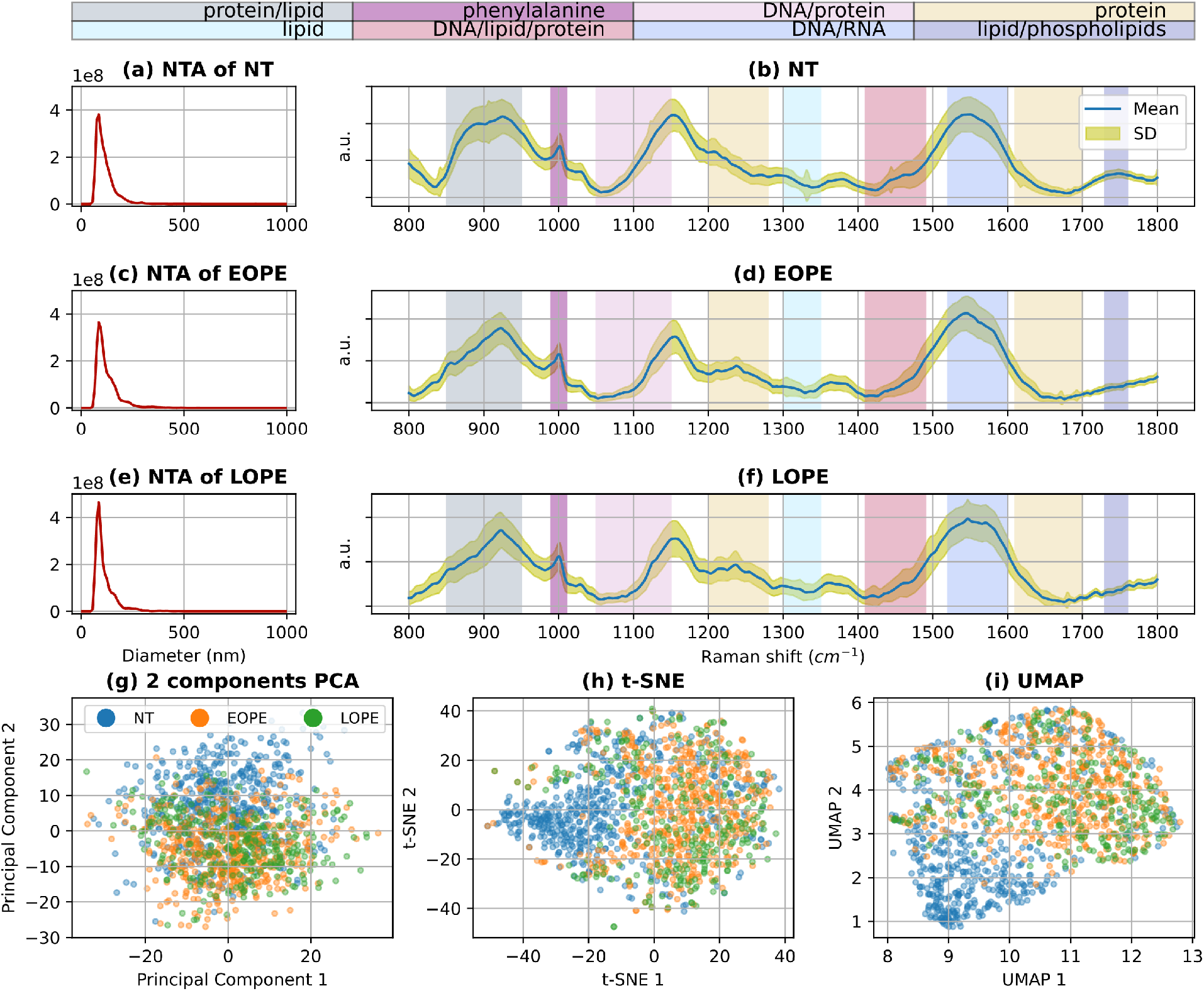
Averaged placental EV data showing NTA size distributions, and normalized SERS spectra for (a-b) NT, (c-d) EOPE, and (e-f) LOPE placental EVs. Distributions of all obtained spectra in two dimensional (g) PCA, (h) t-SNE, and (i) UMAP planes.

Given EVs important and established role in cellular signaling, energy storage, and building of cellular membranes, as well as their clinical association with the vascular wall pathologies, the clear differences in lipid and phospholipid-associated peaks should not be understated. For example, Omatsu *et al*. demonstrated that injecting phosphatidylserine-phosphatidylcholine artificial micro-vesicles induced a preeclampsia-like disease in mice,^65^ while He *et al*., characterized the maternal blood lipidome and demonstrated that phospholipids including phosphadidyl cholines, phosphatidylethanolamines, and ceramides are possible biomarkers for preeclampsia. ^66^ In addition, one previous study used conventional Raman spectroscopy for to characterize maternal serum and found minor but distinct differences in several peaks. ^67^ Confirming the importance of differences in the lipid content of EVs from preeclampsia, Chen *et al*., have recently published that placental EVs that have vesicle-surface exposed phospatidyl serine (identified by annexin V binding) are increased in preeclamptic pregnancies.^68^ However, the exact role of placental EV-lipid content and its role in normotensive and preeclamptic pregnancies have not been fully elucidated to date. These EV SERS findings encourage further lipidomic analyses of these types of EVs to better understand the distinct differences that may be present, and if they could be used as biomarkers outside of SERS analyses.

For the sake of further visualizing of all acquired spectra, PCA, t-SNE, and UMAP were used as dimension reduction visualization techniques to embed the high-dimensional preprocessed spectroscopy data into a lower dimension space as seen in Figure 4g-i. As can be seen in the PCA figure, there is no clear boundary which can separate the NT EV samples from EOPE and LOPE. This is likely due to the fact that PCA transformation preserves the maximum variance of the data in its lower dimension embedding space and is heavily influenced by the inherent heterogeneity of EVs. As these EVs are produced using explant cultures, they are not from a single cell type and should contain diverse biomolecular cargo, ultimately producing diverse Raman spectra.

In contrast, t-SNE and UMAP offer a more clear boundary between NT and preeclamptic EV data. This finding is in agreement with our previously reported use of nonlinear manifold learning for the purpose of the EV Raman data visualization. ^21^ However, due to the inherent heterogeneity of EVs, particularly those produced from mixed cell populations, an improved unsupervised technique was needed to clearly demonstrate the difference between NT, EOPE, and LOPE samples as well as the heterogeneity within individual samples. To accomplish this, spatial K-mean clustering was performed for all of the samples’ normalized spectra as shown in Figure 5a. K-mean clustering investigates the abundance of different types of Raman spectra in each sample by automatically labeling the samples based on their shapes and peaks. The center of each of the clusters are also shown in Fig. 5b.

**Figure 5:**
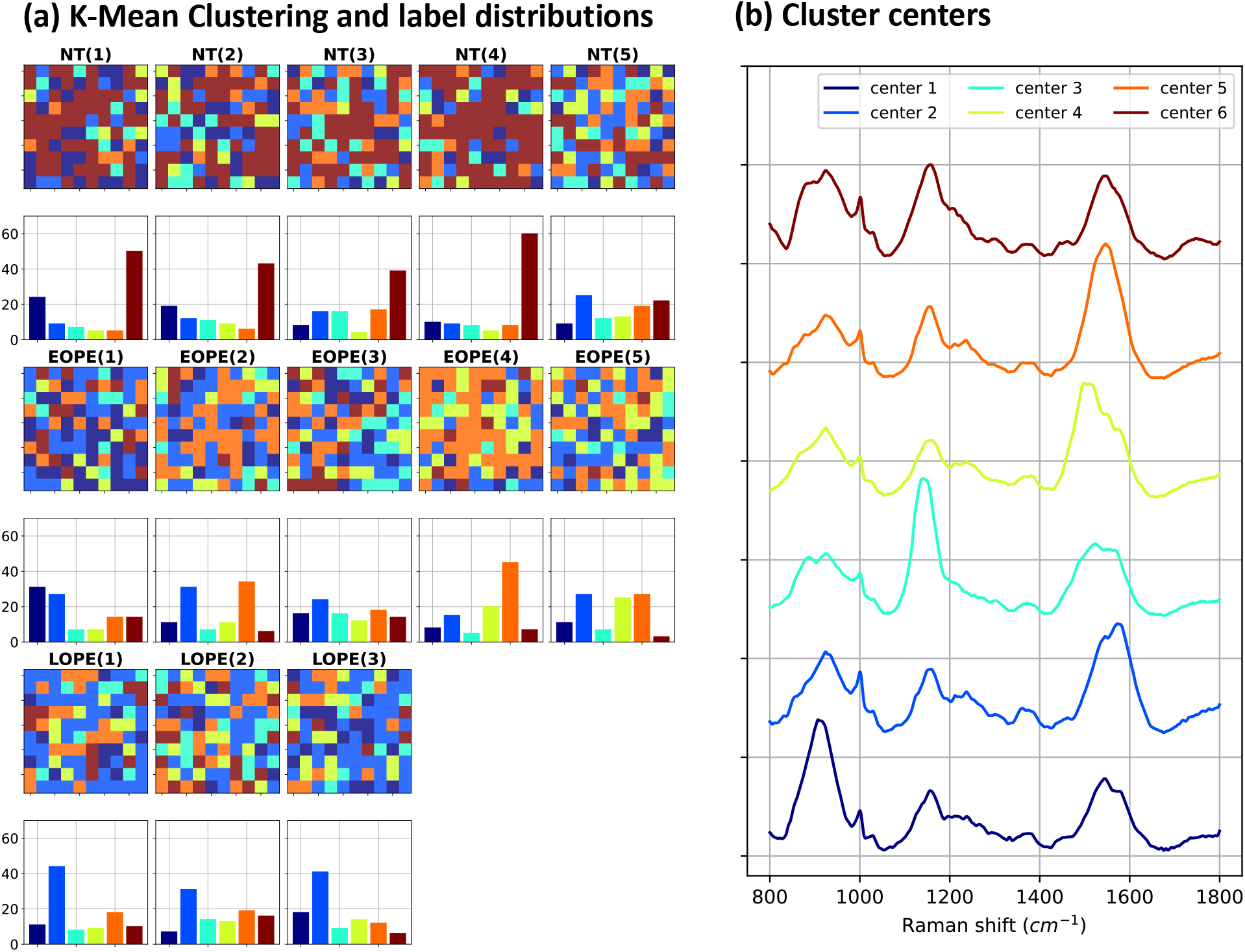
K-mean cluster analysis of LINST SERS using normotensive and preeclamptic EVs from tissue explant cultures showing (a) distribution of K-mean clusters of each of the NT, EOPE and LOPE samples and (b) center of each cluster depicted in (a) with their corresponding colors.

As seen in Fig. 5, there are major differences in the distribution of cluster centers between the NT, EOPE, and LOPE EV samples, implying that there are different mixtures of EV populations in each sample. For instance, center-6 (brown) dominates the NT samples while center-5 (orange) is more common in EOPE samples, and center-2 (light blue) is the most common spectra in the LOPE samples. Fig. 5 also demonstrates the spatial heterogeneity which can exist even from a single explant culture. This is due to the fact that different EVs are positioned at different points on the SERS substrate and thus acquiring spectra across a suitably sized surface is necessary to properly characterize the EV distribution of a given sample.

### EV Classification

Two advanced machine learning methods were employed to classify the placental EV SERS spectra using neural networks. In the first method, a hybrid autoencoder-inspired architecture was used to first reduce the dimensionality of the data to one using dense layers with linear activation. Then, nonlinear activated dense layers were used to classify the samples, as shown in Figure6a. This was done in continuation of the efforts in^69^ and^70^ for the purpose of simultaneous visualization and classification. The second method is based on a deep convolutional neural network for performing the direct classification over the raw spectra with no preprocessing using our previously developed method.^71^

**Figure 6:**
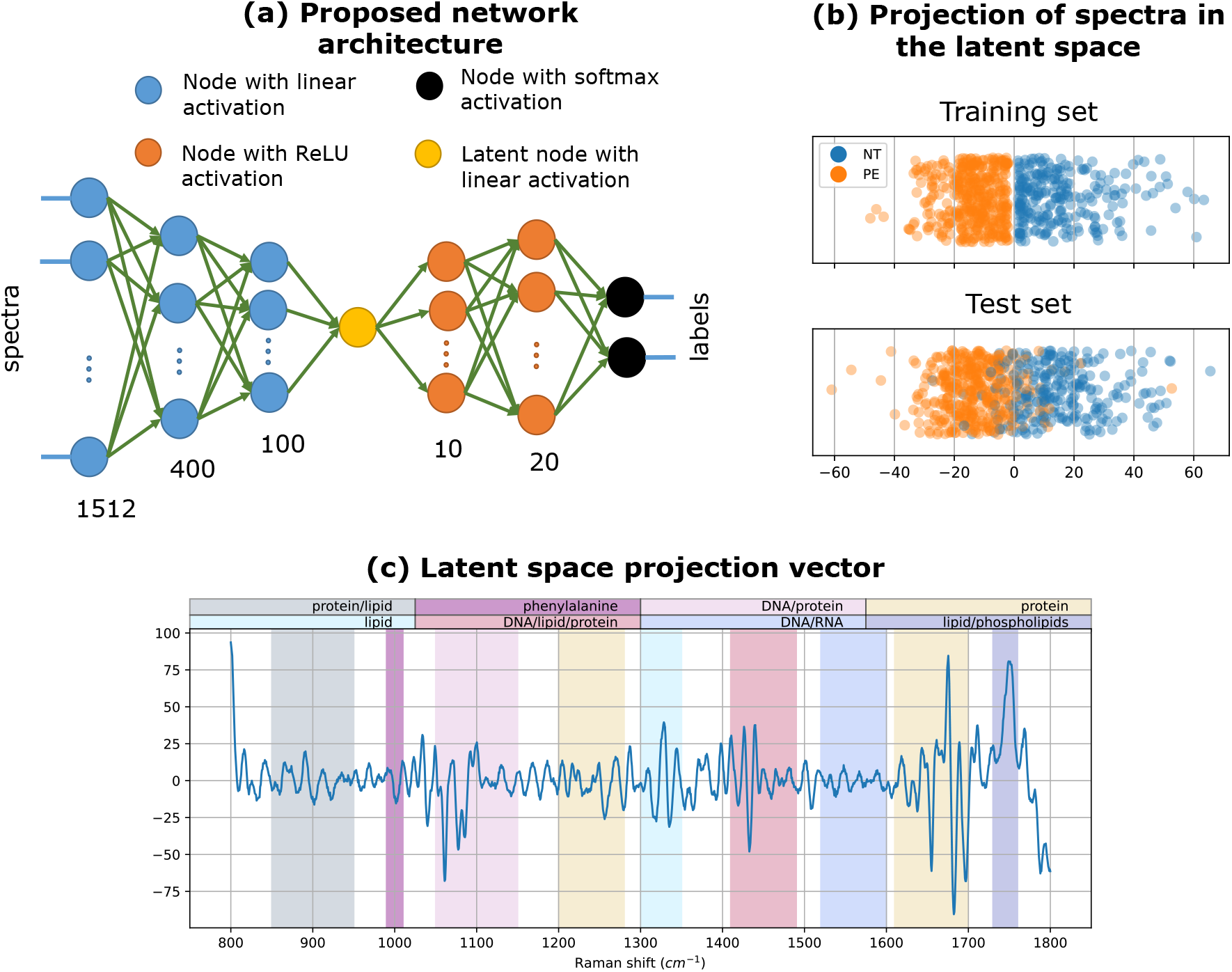
Proposed hybrid autoencoder-inspired classifier showing (a) proposed network design, (b) projection of training and test set in the one dimensional latent space, and (c) calculated direction obtained by the linear activated layers of the network.

#### Bottleneck Classifier and Linear Dimension Reduction

In Figure 6, the first linear dense layers seek a particular direction in the euclidean hyperspace of the data which optimally separates it based on its labels. The main advantage of using this type of network is that the latent space is explainable, such that any obtained values for each of the spectra directly indicates the the presence or absence of specific Raman spectral features. The distribution of the training and test spectra (when 50 percent of the data is randomly chosen for testing purposes) is shown in Figure 6b. As shown, the obtained metric from the training set is a good way to separate the data based on their labels for both training and test sets, as the topology of the training data is recreated with very good precision by the test set. It is important to note that the obtained distribution of the spectra in the latent space is a result of linear transformation, similar to the PCA transformation in figure4. However, while PCA aims to keep the maximum variance of the data in its latent space, the projection in Figure 6b aims to effectively separate them based on their specific labels within the latent space. Another important advantage of this type of classification is its ability to avoid potential issues caused by EV heterogeneity. Similar to autoencoder, the presented network is forced to compress the data and thus rejects randomness in the spectra as much as possible. This important ability was investigated in detail in a similar network for image denoising and compression.^72,73^

The calculated direction using the first linear layers is presented in the Figure 6c. Importantly, as all the first layers are linearly activated, one can easily find the corresponding value of any of the spectra by calculating the dot product of the spectra and the obtained direction and adding the equivalent bias. The obtained result is then classified as the NT if the calculated value is positive and PE if the value is negative. Interestingly, this obtained direction has two distinct positive peaks around 1330 *cm*^−1^ and 1745 *cm*^−1^, which agrees with Figure 4, indicating that lipids or phospholipids are projected onto the positive side of the latent axis in Figure 6b. Finally, this custom network achieved more than 92 percent of accuracy of classification between NT and PE.

#### Deep Convolutional Neural Network

Automatic classification was also carried out using our previously developed deep convolutional neural network over the raw spectral data. This network was designed and optimized specifically for EV SERS spectra and hence leads to much better accuracy over other machine learning methods that are trained using preprocessed data. ^71^ Classification using this network was not done based on the data’s baselines or any randomness such as noise or disturbances, and its sensitivity over spectral shifts indicates the classification is based on the spectral peaks’ positions. To train this network, we first increased the size of the data set for each of the labels (NT and PE) to 5000 spectra using our established data augmentation technique. The augmentation was performed by adding extra Gaussian noise, changing the baselines, small spectral shifts, and finally linear combinations of the spectra within the same label. Then, this network was trained using Adam optimizer as a variant of stochastic gradient descending with learning rate of 10^−4^. Classification accuracy of 96 percent was achieved using this technique, which is significantly higher than the previous method. However this network, like many deep learning algorithms, classifies samples based on characteristics that may not be easily attributed to basic spectral features, such as a single peak’s magnitude. Despite its improved efficiency, this approach is unlike most established diagnostic approaches, where classification based on the abundance known biomolecules is preferred because it can be directly related to known causes or processes of a specific disease. Compared to conventional machine learning classification algorithms such as linear discriminant analysis (LDA), support vector machine (SVM), Random forest (RF), Gaussian process classifier (GPC), or k-nearest neighbors (KNN), the methods proposed above are more robust or more efficient classification approaches (Supplementary Figure 2).

## Conclusions

In this study, we established a simple, tunable, and scalable method of nanoplasmonic sensor fabrication known as laser-induced nanostructuring of SERS-active thin films (LINST). The resulting versatile SERS substrate is suitable for both chemical and EV analyses, as it produces both concave and convex curvature on the surface and redeposits gold nanoparticles of varying size as a byproduct of the laser machining process. To demonstrate its potential in biomedical applications, EVs were cultured and isolated from tissue explant cultures of both normotensive and preeclamptic placentae. Then, using the LINST substrate for EV SERS analysis, they were found to produce classifiably distinct Raman spectra using specifically designed machine learning algorithms.

Our reported results, combined with the abundance and availability of placental EVs within maternal circulation and the power of advanced machine learning classification should encourage further exploration of placenta-derived EV lipidomic analyses, as well as the use of EV SERS for preeclampsia diagnostic or monitoring applications. While this may be possible using the total EV population within the maternal blood, a more specific approach could also be possible by isolating the placenta-specific EVs based on known surface antigens, such as placental alkaline phosphatase or other antigens such as trophoblast glycoprotein/5T4. This could be done either directly on the SERS substrate, which would require compensation for the ligand-specific Raman spectral contribution, or prior to SERS analysis using microfluidic or other EV-capture systems, followed by elution or release.^74–76^

Finally we note that one additional benefit of LINST is that it can be adapted for the nanopatterning of any type of thin film by tuning the laser parameters for the specific target material, and furthermore, it can be done in a single fabrication step. This avoids the need of additional materials or chemicals, or the use of cleanroom environments since the process is performed in an open air atmosphere. Although we have only demonstrated it using a low repetition rate laser, using higher powers and repetition rates along with faster scanning speeds could allow the fabrication times to be reduced by even 100 fold compared to the time required in this preliminary study.^77^

## Materials and Methods

### Thin Film Deposition

Two methods of gold thin film deposition were used to assess their potential for LINST patterning. Magnetron sputter coating with a Q150R sputter coater (Quorum) was used in order to deposit a 200 nm layer of pure gold on the surface of Silicon wafers (p-doped [111]). The process was run with a current of 50 mA which resulted in deposition rate of 6 nm per minute. For the second method, a thermal evaporator was used to firstly deposit 10 nm of chromium followed by 200 nm of gold. The film thickness was measured simultaneously using film thickness monitor (FTM) crystal.

### Laser-Induced Nanostructuring of SERS-Active Thin Films (LINST)

Femtosecond laser surface patterning was conducted on the deposited gold thin films. The laser setup consisted of a Ti:Sapphire laser system combining an ultrafast oscillator (VI-TARA) with a LEGEND regenerative amplifier (Coherent). This system generates 140 fs pulses at a central wavelength of 800 nm and a maximum pulse repetition rate of 1 kHz which was maintained constant in all experiments. At 1 kHz, the laser system the maximum pulse energy was 1.5 mJ and was reduced using a variable attenuator and neutral density filters. The 2 mm laser beam was focused on to the gold surfaces by a 5x Mitutoyo objective with a numerical aperture of 0.14 to achieve a theoretical focused beam waist of 11.2 *μ*m.

Aiming to slowly nanostructure the surface and avoid the complete ablation of the SERS-active thin films, the power was carefully tuned and optimized to test the effect of fluences ranging from 0.05 J/cm^2^ to 0.6 J/cm^2^, and scanning speeds ranging from 0.5 to 1.5 mm/s. The laser’s polarization was set parallel to the scanning direction in all the experiments. The use of these extremely low fluences led to an effective spot size of approximately 2.5 *μ*m (measured from SEM images) since only a very small part of the focused Gaussian beam was above the ablation threshold of the material. It is also important to note that because laser ablation threshold is a highly nonlinear process, the effective beam waist is much smaller than the usual 1/*e* criterion used in the linear regime. For fluences higher than 0.5 J/cm^2^, all the tested scanning speeds resulted in complete gold thin film ablation in certain areas and were therefore discarded as a viable SERS platform option.

Samples were moved using a three-axis motorized stage at different scanning velocities with a position accuracy of 1 *μ*m while the whole process was monitored using an CCD camera. Careful adjustments to the separation of the scanned lines was necessary to ensure that the LINST pattern was homogenuous across the entire substrate. Following the laser nanopatterning, the samples were briefly cleaned with compressed air and kept in a clean atmosphere to prevent any contamination present prior to the SERS experiments.

For smaller fluences, leading to soft LINST (i.e. gold thin-film ablation smaller than thin-film thickness), the effective spot size and line width were characterized via SEM and 1×1.5 mm machined areas were fabricated by adjusting the line spacing according to the measured effective spot size. The fabricated samples were then subjected to Raman chemical test with R6G dye and the structure fabricated with a fluence of 0.2 J/cm^2^, scanning speed of 1.125 mm/s, and a separation between the scanned lines of 2.5 *μ*m showed the best SERS signal enhancement for both types of deposited gold thin films. Therefore, these parameters were used to create extensive identical samples to be used for the chemical and EV SERS testing in this work.

### LINST Surface Characterization

The surface morphology and topography of the LINST samples were visualized using SEM (SU-70, Hitachi) and AFM (Cypher-ES, Asylum Instruments) with a standard ’TAP-150AlG’ cantilever. The AFM scans were performed with a drive frequency of 141.7kHz and 3V free amplitude. All the images were taken in repulsive mode with a set point of 2V.

### EV Isolation from Placentae

Chorionic villi of term placenta were excised into small pieces of approximately 400 mg each for culture and EV isolation (Supplementary Figure 3). Each piece was incubated overnight in a Netwell™ insert in a Falcon® 12-well plate filled with Advanced DMEM/F-12 (Dulbecco’s Modified Eagle Medium/Ham’s F-12, Thermo Fisher Scientific). The media was supplemented with 2% FBS (fetal bovine serum, Life Technologies) and 1% Penicillin-Streptomycin (Life Technologies).

EVs were isolated from the conditioned culture media by differential centrifugation. First, the culture media was centrifuged at 2,000 ×*g* for 5 minutes to pellet any placental debris, cell components, and larger EVs. The resulting supernatant was centrifuged at 20,000 ×*g* for 1 hour at 4°C (Sorvall WX 100+ Ultracentrifuge, ThermoFisher) to pellet large EVs and the remaining supernatant was again centrifuged at 100,000 ×*g* for 1 hour at 4°C to pellet crude small EVs. The small EVs were resuspended in 500 μL of 1x PBS to subsequently perform size exclusion chromatography (SEC) in a 35 nm qEV Original column (Izon) by initially collecting fractions 7-20.

### EV Characterization

EV-rich fractions (7-10) from SEC were initially determined by protein (Pierce™ BCA Protein Assay, ThermoFisher) and particle quantification using nanoparticle tracking analysis (NTA) using an NS300 Nanosight (Malvern Panalytical), then pooled for further analysis. Pooled EVs were diluted at a 1:500 ratio in PBS and three 30 second videos were taken under low flow conditions (Screen gain:1, Camera level:14) and characterized using the Nanosight 3.4 software (Screen gain:10, Detection threshold:6) to calculate mean and mode particle diameters, concentration, and size distributions.

Prior to SERS, EV samples were transferred from PBS buffer to ultrapure water by loading 200 *μ*l of the purified EVs into a Vivaspin 500 (Sartorius AG) centrifugal concentrator with a 100 kDa cutoff and centrifuging at 10,000 ×*g* until most of the PBS had flowed through the filter (roughly 10 minutes). 450 *μ*l of ultrapure water was then added to the filter and the centrifugation process was repeated twice more before finally suspending the EVs in 100 *μ*l ultrapure water.

### Raman Analysis and Machine Learning

Raman spectra were acquired using a LabRAM HR Evolution confocal Raman microscope (Horiba) with either 532 nm or 785 nm excitation wavelengths, depending on the experiment. Microscope objectives of 5× (NA=0.14), 10× (NA=0.26), and 50× (NA=0.42) were used for the different purposes of this work. The laser power was controlled using a neutral density filter and set to 1% to 25% percent of the maximum power (100 mW) depending on the sample test. All measurements were obtained with a 0.1 s to 10 s acquisition time, depending on the laser setup and choice of microscope object with only 1 accumulation. The system utilizes a 600 gr/mm (750) blazed grating in conjunction with a notch filter to remove Rayleigh scattered light. While detection is conducted through a liquid nitrogen (−70 °C) CCD array (1024 × 256 pixels) detector. A confocal hole was set to 250*μ*m in back-scattering geometry. Then, the baseline was established and noise was removed using asymmetric least squares smoothing established in.^78^

### Multivariate Analysis and Machine Learning

PCA and t-SNE transformations and K-mean clustering were carried out using established Sklearn package in python.^79^ All the machine learning and deep learning methods were implemented using the TensorFlow python package.^80^

### Ethics

All placentae were obtained following written informed consent. This study was approved by the Northern A Health and Disability Ethics Committee, approval number NTX/12/06/057/AM1.

## Author Contributions

MK: conceptualization, methodology, software, validation, investigation, writing, MM: conceptualization, methodology, validation, investigation, writing, CA: methodology, resources, SYP: investigation, writing, MML: investigation, WX: supervision, project administration, LWC: conceptualization, writing, funding acquisition, project administration, resources, NGRB: resources, supervision, project administration, funding acquisition, CLH: conceptualization, methodology, validation, investigation, resources, writing, supervision, project administration.

## Acknowledgements

The authors would like to thank the Photon Factory and the Hub for Extracellular Vesicle Investigations at the University of Auckland, and the LEGACY postdoctoral scholars program at The Ohio State University for their continued support. This work was funded by The Health Research Council of New Zealand, Auckland Medical Research Foundation, and the Goodfellow Fund for the Support of Urological Research.

## Graphical TOC Entry

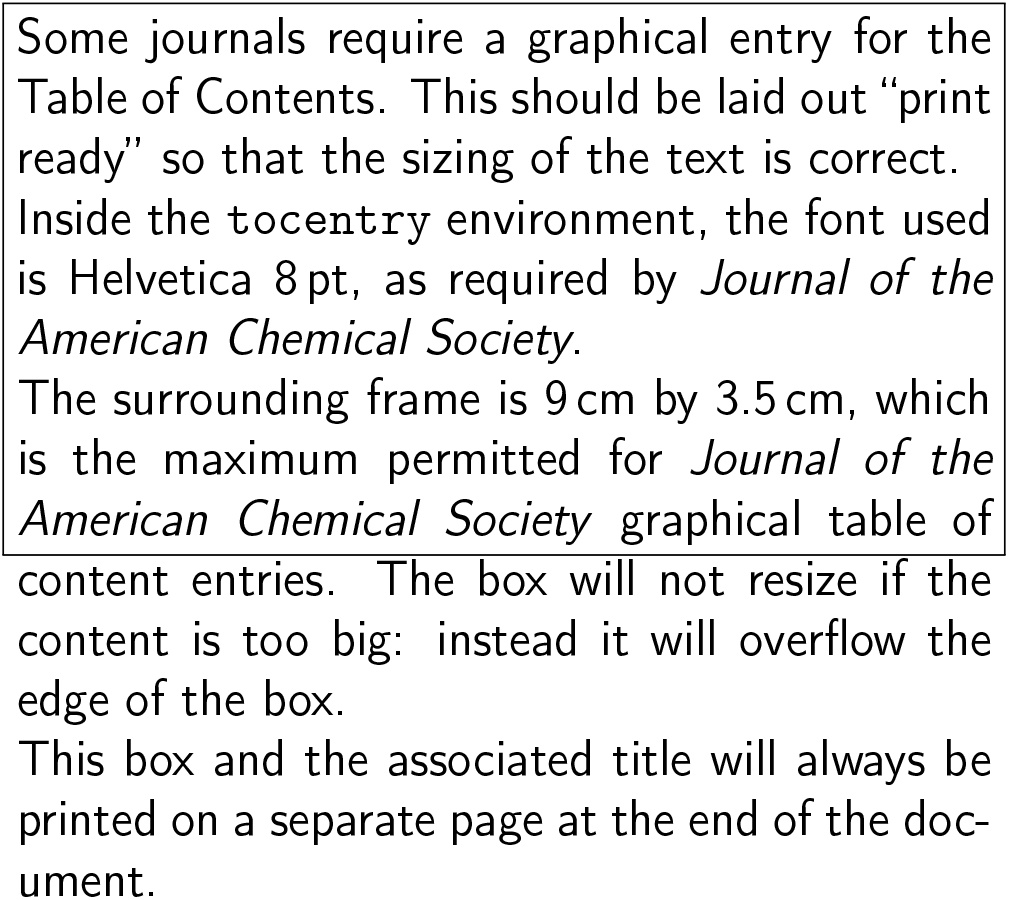

