## Supplementary Figures 1-3 for "Classification of Preeclamptic Placental Extracellular Vesicles Using Femtosecond Laser-fabricated Nanoplasmonic Sensors and Machine Learning"

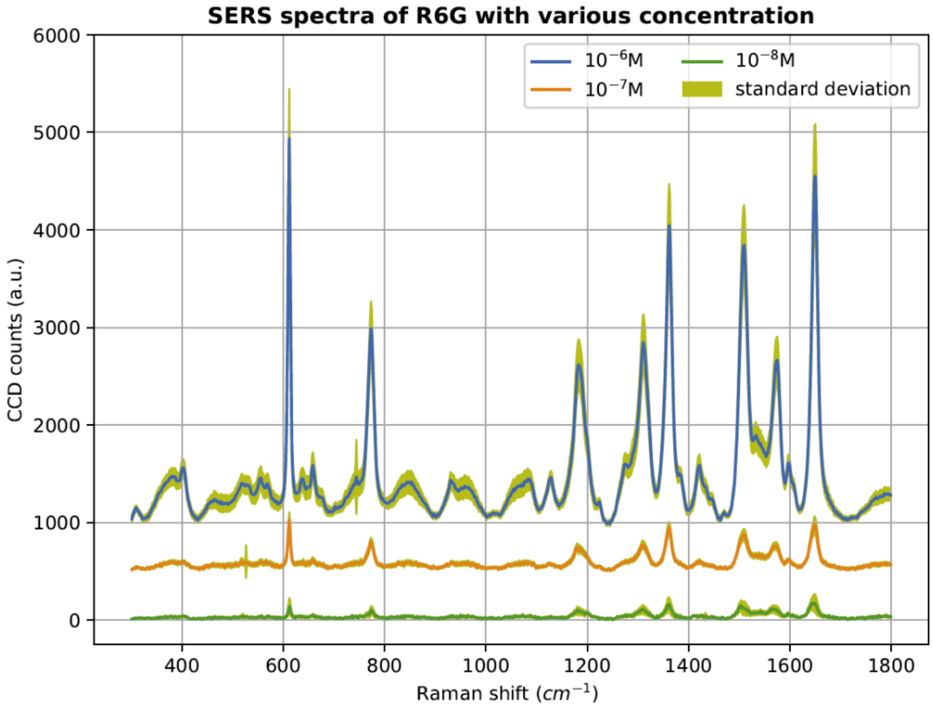


**Supplementary Figure 1.** Raman spectra of R6G with various concentrations obtained using 532nm laser and 1 second acquisition time.

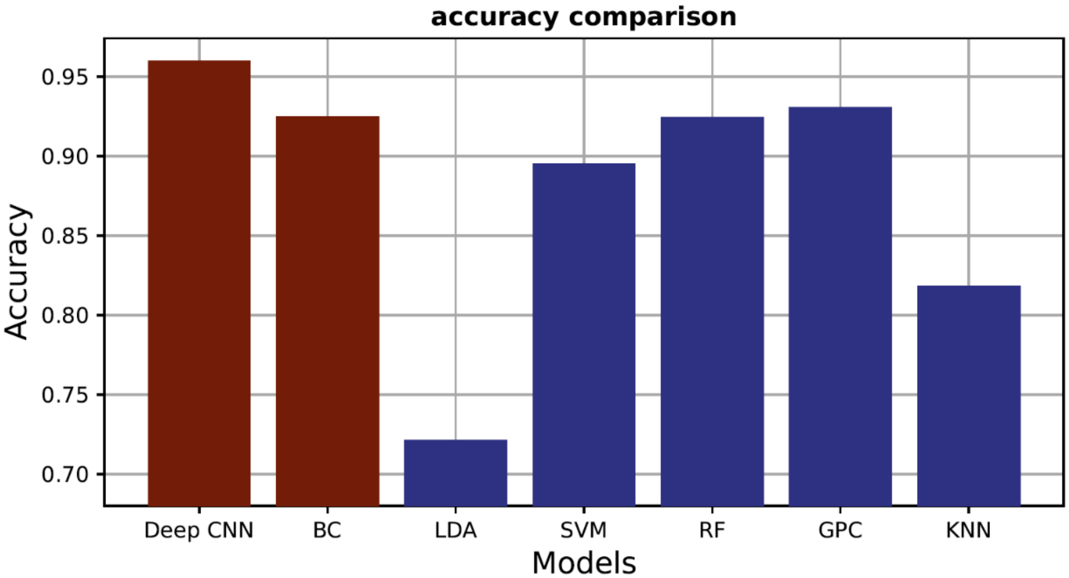


**Supplementary Figure 2.** Comparison of our custom Deep Convolutional Neural Network (CNN) and Bottleneck Classifier (BC) algorithms with conventional machine learning classifiers including Linear Discriminant Analysis (LDA), Support Vector Machine (SVM), Random Forest (RF), Gaussian Process Classifier (GPC), and K-nearest Neighbor (KNN).

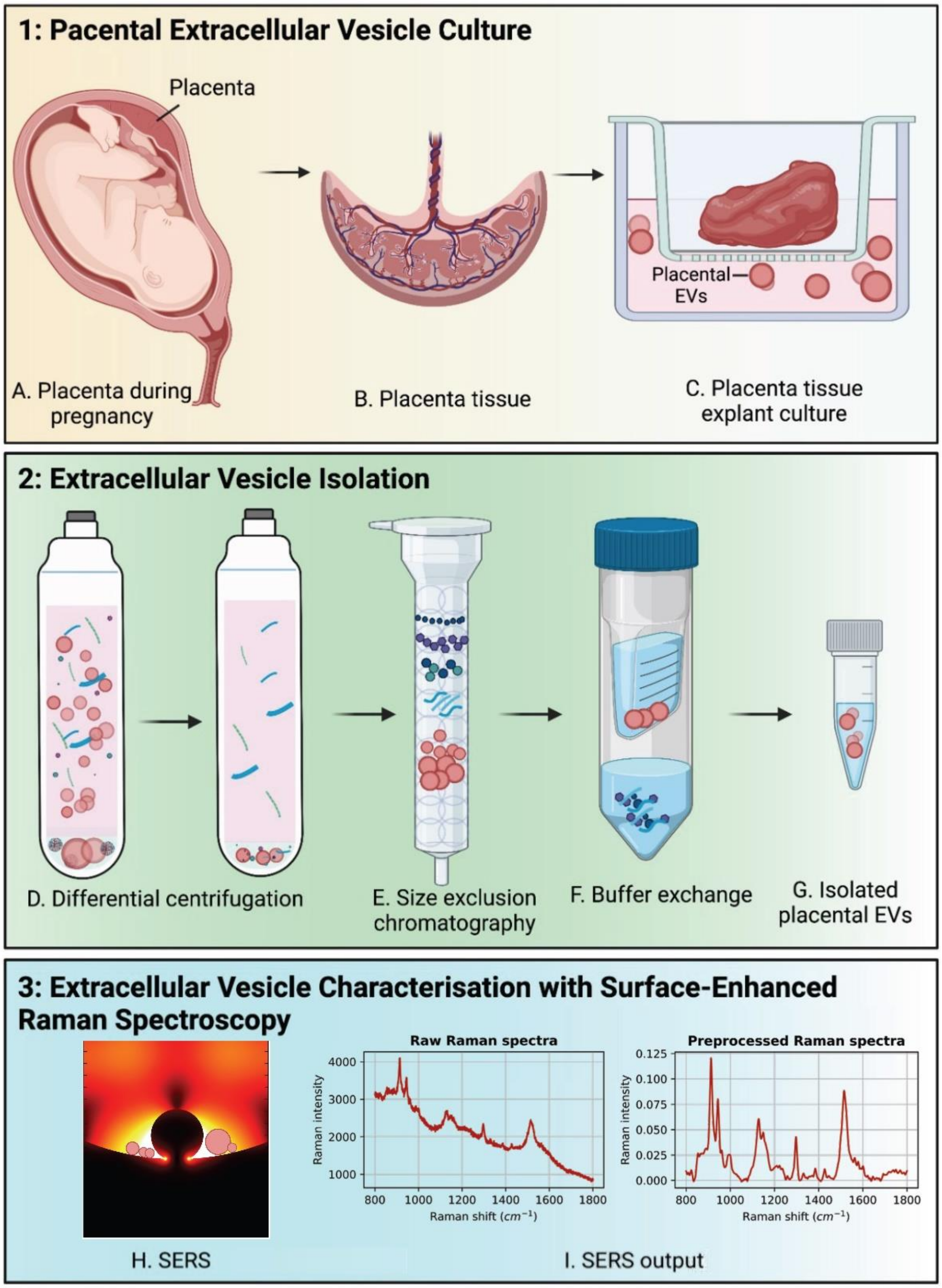


**Supplementary Figure 3.** Placental explant culture and EV isolation schematic showing (A-C) isolation of explants from term placenta and culture in Netwell™ inserts, (D-G) EV isolation using differential ultracentrifugation, size exclusion chromatography, and ultrafiltration for buffer exchange immediately prior to SERS, and (H-I) EV characterization using SERS.
